# Evaluation of the External RNA Controls Consortium (ERCC) reference material using a modified Latin square design

**DOI:** 10.1101/034868

**Authors:** PS Pine, SA Munro, JR Parsons, J McDaniel, Lucas A Bergstrom, J Lozach, TG Myers, Q Su, SM Jacobs-Helber, M Salit

**Affiliations:** Joint Initiative for Metrology in Biology, National Institute of Standards and Technology, Stanford, CA, 94305, USA; Genomics Research and Development, Agilent Technologies, Santa Clara, CA, 95051, USA; Illumina, Inc., San Diego, CA, 92122, USA; National Institute of Allergy and Infectious Diseases, Bethesda, MD, 20892, USA; AIBioTech, Inc., Richmond, VA, 23235, USA

**Keywords:** ERCC, gene expression, microarray, RNA controls, RNA sequencing, RNA-Seq, spike-in controls

## Abstract

**BACKGROUND:** Highly multiplexed assays for quantitation of RNA transcripts are being used in many areas of biology and medicine. Using data generated by these transcriptomic assays requires measurement assurance with appropriate controls. Methods to prototype and evaluate multiple RNA controls were developed as part of the External RNA Controls Consortium (ERCC) assessment process. These approaches included a modified Latin square design to provide a broad dynamic range of relative abundance with known differences between four complex pools of ERCC RNA transcripts spiked into a human liver total RNA background.

**RESULTS:** ERCC pools were analyzed on four different microarray platforms: Agilent 1- and 2-color, Illumina bead, and NIAID lab-made spotted microarrays; and two different second-generation sequencing platforms: the Life Technologies 5500xl and the Illumina HiSeq 2500. Individual ERCCs were assessed for reproducible performance in signal response to concentration among the platforms. Most demonstrated linear behavior if they were not located near one of the extremes of the dynamic range. Performance issues with any individual ERCC transcript could be attributed to detection limitations, platform-specific target probe issues, or potential mixing errors. Collectively, these pools of spike-in RNA controls were evaluated for suitability as surrogates for endogenous transcripts to interrogate the performance of the RNA measurement process of each platform. The controls were useful for establishing the dynamic range of the assay, as well as delineating the useable region of that range where differential expression measurements, expressed as ratios, would be expected to be accurate.

**CONCLUSIONS:** The modified Latin square design presented here uses a composite testing scheme for the evaluation of multiple performance characteristics: linear performance of individual controls, signal response within dynamic range pools of controls, and ratio detection between pairs of dynamic range pools. This compact design provides an economical sample format for the evaluation of multiple external RNA controls within a single experiment per platform. These results indicate that well-designed pools of RNA controls, spiked-into samples, provide measurement assurance for endogenous gene expression experiments.

## Background

In 2003, the National Institute of Standards and Technology (NIST) hosted a meeting to discuss the need for a universal RNA reference material, which could be used for gene expression profiling assays [1]. As a result of this effort, the External RNA Controls Consortium (ERCC) was formed, of which NIST is a founding member and host. The ERCC assembled a sequence library of 176 DNA sequences that could be transcribed into RNA to serve as controls in systems used to measure gene expression [2, 3]. These controls were cataloged as ERCC-00001 through ERCC-00176, and are collectively referred to as ERCCs in this manuscript. These were evaluated and a subset was selected for dissemination as a standard. A set of 96 controls are now available as a set of sequence-certified DNA plasmids, NIST Standard Reference Material (SRM) 2374 [4].

In the final phase of evaluation, an experimental design for assessing the combined performance of ERCCs prepared as complex RNA pools was used. Each ERCC subpool was designed to have a 2^20^ dynamic range of abundance of controls, and particular controls in the different pools were present in different abundances according to a modified Latin square design. This design provides known relative differences between the pools across a large dynamic range of abundance (**Figure 1**). With this design, individual ERCCs were assessed for their signal response to 1.5-, 2.5-, and 4-fold increases in concentration. Pairwise comparisons of these pools also provides for an assessment of ratio-based performance as a function of dynamic range. Initially assessed with three different microarray platforms, these same pools were subsequently measured by RNA sequencing (RNA-Seq) with two second-generation (NGS) sequencing platforms. The data from these two sets of experiments, corresponding to the 96 controls of the SRM, are presented here.

**Figure 1.**
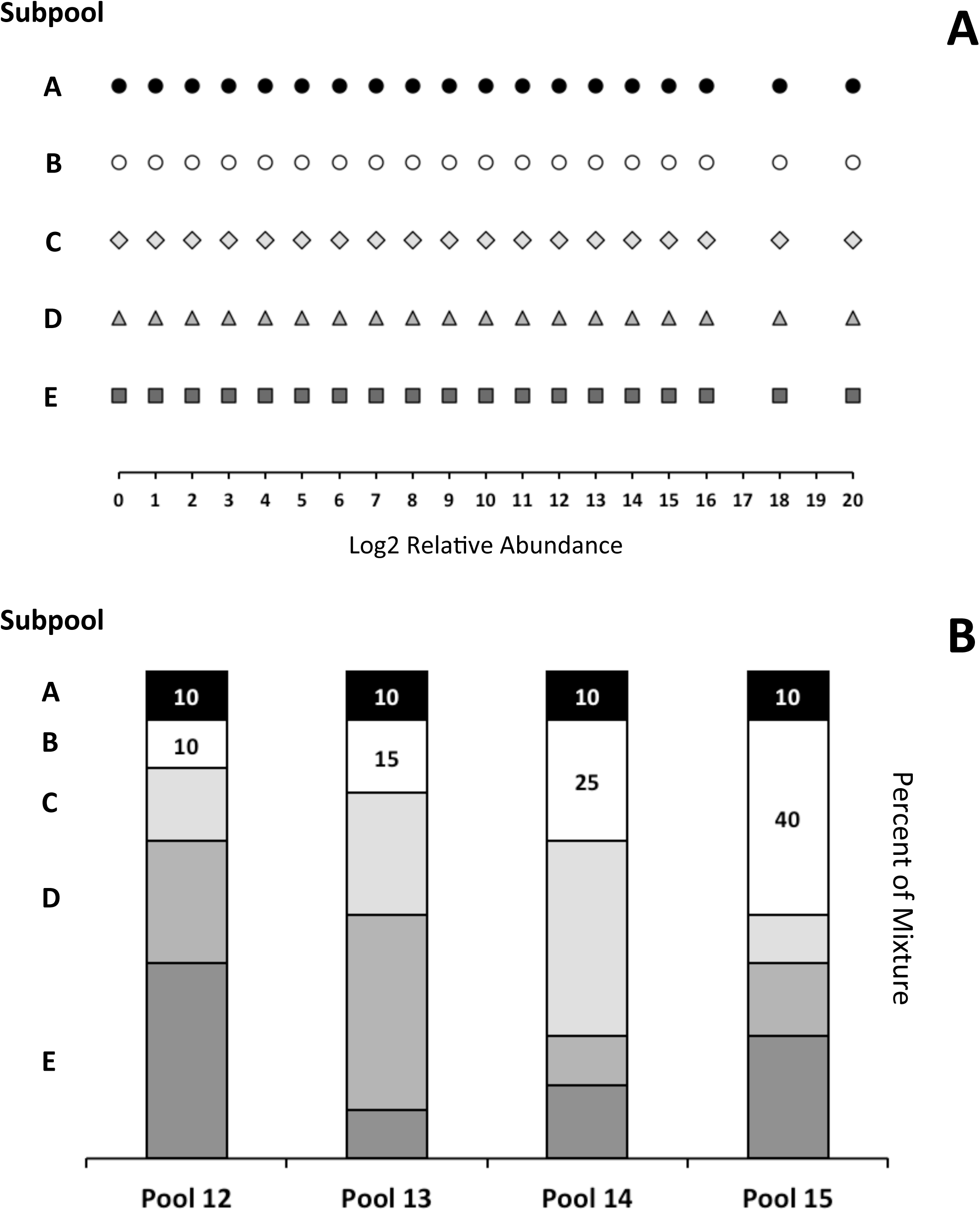
*Latin square plus pool design*. Panel A shows a schematic design of the relative abundance of 95 unique ERCC distributed into 5 subpools. Panel B shows the proportion of each subpool within each pool. Subpools B – E are mixed using a Latin square of proportions 40, 25, 15, and 10 percent, plus subpool A as an additional 10 percent component of each. Subpools A, B, C, D, and E are shaded, black, white, light grey, medium grey, and dark grey, respectively. Refer to Table 1 for the target relative abundance of an ERCC used in the design of each pool.

**Table 1.**
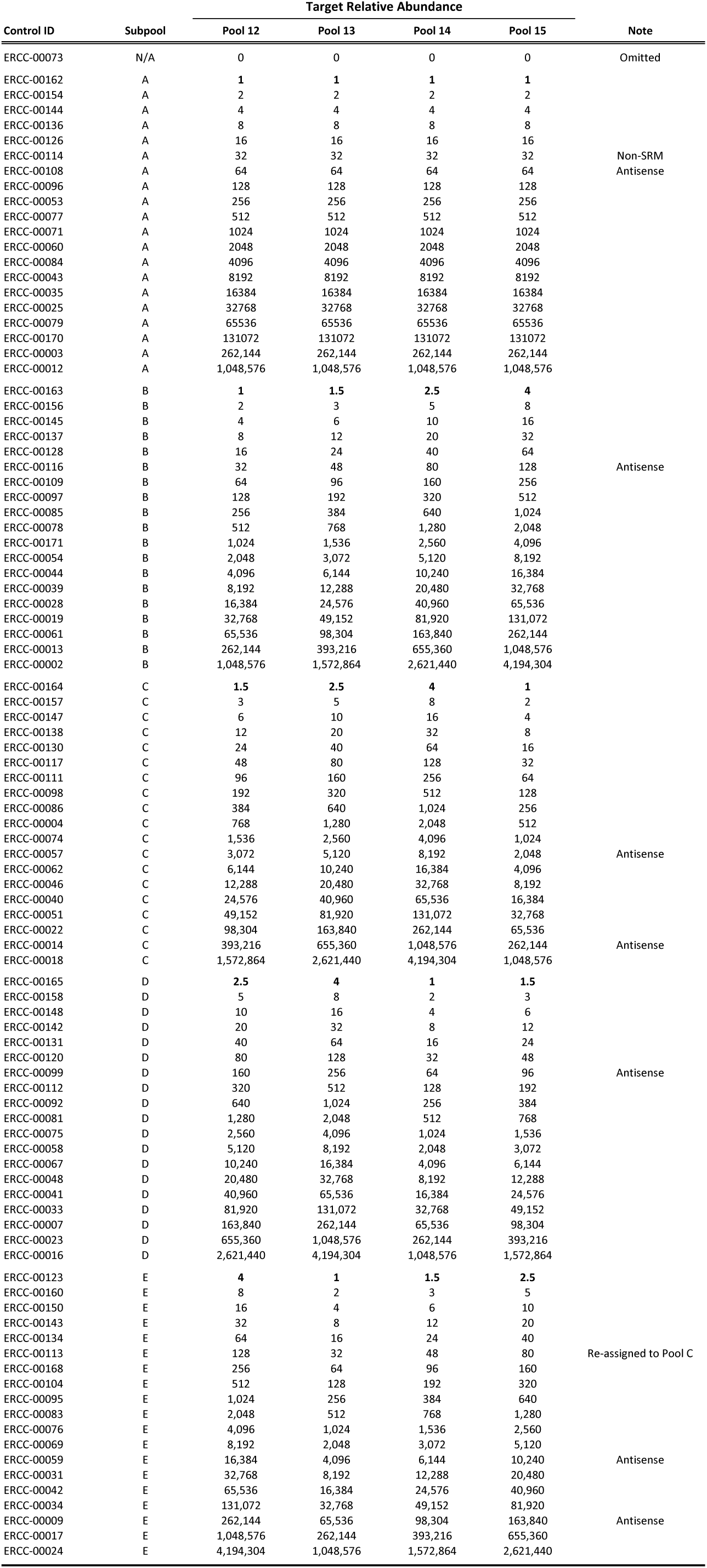
Distribution of ERCCs among pools and mixtures.

| Control ID | Subpool | Target Relative Abundance |  |  |  | Note |
| --- | --- | --- | --- | --- | --- | --- |
|  |  | Pool 12 | Pool 13 | Pool 14 | Pool 15 |  |
| ERCC-00073 | N/A | 0 | 0 | 0 | 0 | Omitted |
| ERCC-00162 | A | 1 | 1 | 1 | 1 |  |
| ERCC-00154 | A | 2 | 2 | 2 | 2 |  |
| ERCC-00144 | A | 4 | 4 | 4 | 4 |  |
| ERCC-00136 | A | 8 | 8 | 8 | 8 |  |
| ERCC-00126 | A | 16 | 16 | 16 | 16 |  |
| ERCC-00114 | A | 32 | 32 | 32 | 32 | Non-SRM |
| ERCC-00108 | A | 64 | 64 | 64 | 64 | Antisense |
| ERCC-00096 | A | 128 | 128 | 128 | 128 |  |
| ERCC-00053 | A | 256 | 256 | 256 | 256 |  |
| ERCC-00077 | A | 512 | 512 | 512 | 512 |  |
| ERCC-00071 | A | 1024 | 1024 | 1024 | 1024 |  |
| ERCC-00060 | A | 2048 | 2048 | 2048 | 2048 |  |
| ERCC-00084 | A | 4096 | 4096 | 4096 | 4096 |  |
| ERCC-00043 | A | 8192 | 8192 | 8192 | 8192 |  |
| ERCC-00035 | A | 16384 | 16384 | 16384 | 16384 |  |
| ERCC-00025 | A | 32768 | 32768 | 32768 | 32768 |  |
| ERCC-00079 | A | 65536 | 65536 | 65536 | 65536 |  |
| ERCC-00170 | A | 131072 | 131072 | 131072 | 131072 |  |
| ERCC-00003 | A | 262,144 | 262,144 | 262,144 | 262,144 |  |
| ERCC-00012 | A | 1,048,576 | 1,048,576 | 1,048,576 | 1,048,576 |  |
| ERCC-00163 | B | 1 | 1.5 | 2.5 | 4 |  |
| ERCC-00156 | B | 2 | 3 | 5 | 8 |  |
| ERCC-00145 | B | 4 | 6 | 10 | 16 |  |
| ERCC-00137 | B | 8 | 12 | 20 | 32 |  |
| ERCC-00128 | B | 16 | 24 | 40 | 64 |  |
| ERCC-00116 | B | 32 | 48 | 80 | 128 | Antisense |
| ERCC-00109 | B | 64 | 96 | 160 | 256 |  |
| ERCC-00097 | B | 128 | 192 | 320 | 512 |  |
| ERCC-00085 | B | 256 | 384 | 640 | 1,024 |  |
| ERCC-00078 | B | 512 | 768 | 1,280 | 2,048 |  |
| ERCC-00171 | B | 1,024 | 1,536 | 2,560 | 4,096 |  |
| ERCC-00054 | B | 2,048 | 3,072 | 5,120 | 8,192 |  |
| ERCC-00044 | B | 4,096 | 6,144 | 10,240 | 16,384 |  |
| ERCC-00039 | B | 8,192 | 12,288 | 20,480 | 32,768 |  |
| ERCC-00028 | B | 16,384 | 24,576 | 40,960 | 65,536 |  |
| ERCC-00019 | B | 32,768 | 49,152 | 81,920 | 131,072 |  |
| ERCC-00061 | B | 65,536 | 98,304 | 163,840 | 262,144 |  |
| ERCC-00013 | B | 262,144 | 393,216 | 655,360 | 1,048,576 |  |
| ERCC-00002 | B | 1,048,576 | 1,572,864 | 2,621,440 | 4,194,304 |  |
| ERCC-00164 | C | 1.5 | 2.5 | 4 | 1 |  |
| ERCC-00157 | C | 3 | 5 | 8 | 2 |  |
| ERCC-00147 | C | 6 | 10 | 16 | 4 |  |
| ERCC-00138 | C | 12 | 20 | 32 | 8 |  |
| ERCC-00130 | C | 24 | 40 | 64 | 16 |  |
| ERCC-00117 | C | 48 | 80 | 128 | 32 |  |
| ERCC-00111 | C | 96 | 160 | 256 | 64 |  |
| ERCC-00098 | C | 192 | 320 | 512 | 128 |  |
| ERCC-00086 | C | 384 | 640 | 1,024 | 256 |  |
| ERCC-00004 | C | 768 | 1,280 | 2,048 | 512 |  |
| ERCC-00074 | C | 1,536 | 2,560 | 4,096 | 1,024 |  |
| ERCC-00057 | C | 3,072 | 5,120 | 8,192 | 2,048 | Antisense |
| ERCC-00062 | C | 6,144 | 10,240 | 16,384 | 4,096 |  |
| ERCC-00046 | C | 12,288 | 20,480 | 32,768 | 8,192 |  |
| ERCC-00040 | C | 24,576 | 40,960 | 65,536 | 16,384 |  |
| ERCC-00051 | C | 49,152 | 81,920 | 131,072 | 32,768 |  |
| ERCC-00022 | C | 98,304 | 163,840 | 262,144 | 65,536 |  |
| ERCC-00014 | C | 393,216 | 655,360 | 1,048,576 | 262,144 | Antisense |
| ERCC-00018 | C | 1,572,864 | 2,621,440 | 4,194,304 | 1,048,576 |  |
| ERCC-00165 | D | 2.5 | 4 | 1 | 1.5 |  |
| ERCC-00158 | D | 5 | 8 | 2 | 3 |  |
| ERCC-00148 | D | 10 | 16 | 4 | 6 |  |
| ERCC-00142 | D | 20 | 32 | 8 | 12 |  |
| ERCC-00131 | D | 40 | 64 | 16 | 24 |  |
| ERCC-00120 | D | 80 | 128 | 32 | 48 |  |
| ERCC-00099 | D | 160 | 256 | 64 | 96 | Antisense |
| ERCC-00112 | D | 320 | 512 | 128 | 192 |  |
| ERCC-00092 | D | 640 | 1,024 | 256 | 384 |  |
| ERCC-00081 | D | 1,280 | 2,048 | 512 | 768 |  |
| ERCC-00075 | D | 2,560 | 4,096 | 1,024 | 1,536 |  |
| ERCC-00058 | D | 5,120 | 8,192 | 2,048 | 3,072 |  |
| ERCC-00067 | D | 10,240 | 16,384 | 4,096 | 6,144 |  |
| ERCC-00048 | D | 20,480 | 32,768 | 8,192 | 12,288 |  |
| ERCC-00041 | D | 40,960 | 65,536 | 16,384 | 24,576 |  |
| ERCC-00033 | D | 81,920 | 131,072 | 32,768 | 49,152 |  |
| ERCC-00007 | D | 163,840 | 262,144 | 65,536 | 98,304 |  |
| ERCC-00023 | D | 655,360 | 1,048,576 | 262,144 | 393,216 |  |
| ERCC-00016 | D | 2,621,440 | 4,194,304 | 1,048,576 | 1,572,864 |  |
| ERCC-00123 | E | 4 | 1 | 1.5 | 2.5 |  |
| ERCC-00160 | E | 8 | 2 | 3 | 5 |  |
| ERCC-00150 | E | 16 | 4 | 6 | 10 |  |
| ERCC-00143 | E | 32 | 8 | 12 | 20 |  |
| ERCC-00134 | E | 64 | 16 | 24 | 40 |  |
| ERCC-00113 | E | 128 | 32 | 48 | 80 | Re-assigned to Pool C |
| ERCC-00168 | E | 256 | 64 | 96 | 160 |  |
| ERCC-00104 | E | 512 | 128 | 192 | 320 |  |
| ERCC-00095 | E | 1,024 | 256 | 384 | 640 |  |
| ERCC-00083 | E | 2,048 | 512 | 768 | 1,280 |  |
| ERCC-00076 | E | 4,096 | 1,024 | 1,536 | 2,560 |  |
| ERCC-00069 | E | 8,192 | 2,048 | 3,072 | 5,120 |  |
| ERCC-00059 | E | 16,384 | 4,096 | 6,144 | 10,240 | Antisense |
| ERCC-00031 | E | 32,768 | 8,192 | 12,288 | 20,480 |  |
| ERCC-00042 | E | 65,536 | 16,384 | 24,576 | 40,960 |  |
| ERCC-00034 | E | 131,072 | 32,768 | 49,152 | 81,920 |  |
| ERCC-00009 | E | 262,144 | 65,536 | 98,304 | 163,840 | Antisense |
| ERCC-00017 | E | 1,048,576 | 262,144 | 393,216 | 655,360 |  |
| ERCC-00024 | E | 4,194,304 | 1,048,576 | 1,572,864 | 2,621,440 |  |

## Methods

### Pool Design

The ERCCs were distributed into 5 subpools (A – E), each containing a unique set of controls (see Fig. 1A). These subpools were prepared at AIBioTech (formerly CBI Services, Richmond, VA) to ERCC specifications. This design results in the relative abundance within each subpool covering a dynamic range of 2^20^. Subpools A – E were then mixed by volume in a modified Latin square design to create 4 different pools (see **Fig. 1B** and **Table 1**). Subpools B – E have different relative abundances between the four pools (in a Latin square design), while subpool A is held at a constant proportion (the “modification”). In addition, the ERCCs in subpools B – E participate in 6 pairwise comparisons between pools to produce ratios of 4-, 2.7-, 2.5-, 1.7-, 1.6, and 1.5-to-1 (Fig. 1B and Suppl. Figs. 1 – 4). The ERCCs in subpool A are always present at 10% in any of the pools, and create the 1-to-1 component in any of the 6 possible pairwise comparisons. These pools were designated as Pools 12, 13, 14, and 15 in the set of pools developed for ERCC testing [2]. Each ERCC of these pools was spiked into a common “background” of human liver total RNA (Ambion) to create 4 corresponding samples. Each microarray test site determined the relative amount of spike-in pools to add to the background. Agilent, Illumina, and NIAID used 0.144%, 0.25%, and 0.265% (wt/wt) of ERCC pool per total liver RNA, respectively. For the sequencing test sites total RNA samples were spiked at NIST at 0.3% (wt/wt) and then sequenced by NIST and Illumina.

The ERCC molecules used in these pools were prepared by *in vitro* transcription of polymerase chain reaction (PCR) products representing “candidate” sequences prior to the release of NIST SRM 2374. The plasmids were designed to produce either “sense” or “antisense” RNA controls [4]. In this study, seven of these ERCC transcripts were determined to be antisense using a stranded RNA-Seq protocol (see **Table 1**) and were excluded from further data analysis, because the microarrays were designed to detect “sense” RNA controls.

### Microarray measurements

#### Samples were measured at each test site using the following methods

The NIAID in-house spotted microarrays contain long (70-mer) oligonucleotides designed to hybridize the ERCC transcripts printed on epoxy-coated glass slides (Corning) in quadruplicate using an OmniGrid robot (Genomic Solutions) with 16 SMP3 print tips (Telechem). RNA was reverse transcribed using Oligo dT primer (12–20 mer) mix (Invitrogen) and Superscript II reverse transcriptase (Invitrogen). Fluorescent Cy-Dye-dUTP (GE) nucleotide was incorporated into first-strand cDNA during the reverse transcription. After degradation of the mRNA template strand, labeled single-stranded cDNA target was purified using Vivaspin 500 (10K, Millipore). Hybridization was performed at 45 C°, for 16 hours on a MAUI hybridization station. The arrays were washed twice in 1X SSC and 0.05% SDS and twice in 0.1X SSC, then air dried. Microarrays were scanned on GenePix 4000B (Axon) at 10 micron resolution. GenePix Pro software was used for image analysis. Median pixel intensity (no background subtraction) was taken for each of the 4 replicate spots, the median of these four values was taken to represent the data.

The Agilent microarrays (8×60K Agilent G3 8-pack format with the Design ID 022439) contain 60-mer oligonucleotide probes synthesized *in situ* onto slides using a proprietary non-contact industrial inkjet printing process. Labeled cRNA for both the one-color and two-color microarray experiments was prepared using the Agilent Low Input Quick Amp Labeling Kit, Two-Color (5190–2306). RNA was reverse transcribed using AffinityScript RT, Oligo(dT) Promoter Primer, and T7 RNA Polymerase. Fluorescent Cy-Dye-dCTP nucleotide was incorporated during cRNA synthesis and amplification. Microarrays were hybridized at 65^o^C for 17 hours. All microarrays were scanned in one batch in random order using default settings for Agilent C Scanner using a single pass over the scan area at a resolution of 3 μm and a 20-bit scan type. Data was extracted with Agilent Feature Extraction Software (ver. 10.7.3.1) using the default settings for either the one-color protocol or the two-color protocol.

The Illumina Human-6 Expression BeadChips contain 50-mer oligonucleotide probes with a 29-mer address sequences attached to beads held in etched microwells. RNA was reverse transcribed using a T7 Oligo(dT) primer containing a T7 promoter sequence. Biotinylated cRNA was prepared using the Illumina TotalPrep RNA Amplification Kit (Ambion). BeadChips were hybridized at 58^o^C for 14 – 20 hours, washed, and labeled with streptavidin-Cy3. BeadChips were scanned with the Illumina iScan System. Intensity values are determined for every bead and summarized for each bead type. For more details refer to the Whole-Genome Gene Expression Direct Hybridization Assay Guide (Illumina, part no. 11322355).

### RNA Sequencing measurements

NIST prepared samples of spiked liver total RNA for sequencing analysis with the 5500xl at NIST and the HiSeq 2500 at Illumina. Prior to library preparation samples were depleted of ribosomal RNA. The 5500xl experiment produced an average of 23,866,495 single-ended reads (75 base) per sample and the HiSeq 2500 experiment yielded an average of 48,168,710 paired-end reads (2 × 75 base). For both platforms sequence reads were aligned against a reference sequence consisting of the human genome (hg19) and the ERCC transcript sequences of SRM 2374 (Note: ERCC-00114 is not part of the SRM and not included as part of the reference transcriptome). Alignment and quantification of sequence reads to obtain per transcript counts was performed with the LifeScope bioinformatic analysis suite (Life Technologies) for 5500×l data and the Tophat-Cufflinks suite was used for HiSeq 2500 data [5,6].

## Results and Discussion

For each of the platforms, if the ERCC spike-in pools are added to the background RNA in the proper proportion, then the 2^20^ range of relative abundance will cover the distribution of the endogenous transcript signals. In the first set of experiments, each microarray platform provider empirically determined in pilot studies their chosen spike-in proportion to add to the total RNA background (not shown). Agilent used 0.144% (wt/wt) for both one-color and two-color arrays, and Illumina and NIAID used 0.25%, and 0.265%, respectively. For the RNA-Seq experiments, ERCC pools were added to the background at NIST at 0.3% and shared with the Illumina site. The LifeTech 5500xl and Illumina HiSeq measurements were performed at NIST and Illumina, respectively. The distribution of ERCC signals relative to the endogenous liver background transcripts are shown for all platforms in **Table 2**. For all sites, the dynamic range of the signals from the controls matched the range of signal expression from the endogenous genes of the liver background. This supports the use of these signals to derive metrics useful for characterizing each measurement system.

**Table 2.** Dynamic Range Coverage.

| Platform | Units | Subset | ERCC-00073 | Minimum | Maximum | Range |
| --- | --- | --- | --- | --- | --- | --- |
| Illumina<br>Bead | log<sub>2</sub> signal | ERCC | 5.81 ± 0.08 | 5.72 ± 0.07 | 13.88 ± 0.11 | 8.16 ± 0.13 |
|  |  | BKGD<sup>1</sup> |  | 5.35 ± 0.07 | 14.34 ± 0.08 | 8.99 ± 0.10 |
| NIAID<br>In-house | log<sub>2</sub> signal | ERCC | 5.53 ± 0.02 | 5.53 ± 0.03 | 15.83 ± 0.38 | 10.30 ± 0.39 |
|  |  | BKGD |  | 5.20 ± 0.01 | 15.26 ± 0.27 | 10.06 ± 0.27 |
| Agilent<br>One-color | log<sub>2</sub> signal | ERCC | 2.62 ± 0.14 | 2.57 ± 0.16 | 20.73 ± 0.18 | 18.16 ± 0.24 |
|  |  | BKGD |  | 2.41 ± 0.10 | 20.66 ± 0.06 | 18.25 ± 0.12 |
| Agilent<br>Two-color | log<sub>2</sub> signal | ERCC | 2.57 ± 0.06 | 2.40 ± 0.06 | 18.37 ± 0.10 | 15.98 ± 0.12 |
|  |  | BKGD |  | 4.40 ± 0.15 | 20.00 ± 0.10 | 15.60 ± 0.19 |
| Illumina<br>HiSeq | log<sub>2</sub> FPKM | ERCC | *undetected* | -4.98 ± 0.67 | 14.58 ± 0.27 | 19.56 ± 0.72 |
|  |  | BKGD |  | -6.34 ± 0.40 | 18.27 ± 0.05 | 24.61 ± 0.40 |
| LifeTech<br>SOLiD | log<sub>2</sub> RPKM<sup>2</sup> | ERCC | *undetected* | -3.26 ± 0.38 | 16.47 ± 0.34 | 19.73 ± 0.51 |
|  |  | BKGD |  | -6.64 ± 0.00 | 17.30 ± 0.35 | 23.94 ± 0.35 |

<sup>1</sup>All transcripts measured in the total human liver RNA background.
<sup>2</sup>Minimum RPKM value reported is truncated at 0.01 for all replicates.

### Dose-response and Outlier Detection

For each platform, we can determine whether the analytical signal (fluorescence intensity in microarrays or length normalized counts in sequencing) changes with the concentration of an analyte (the ERCC being measured). For each control, the signal from each pool can be plotted against the corresponding relative abundance (**Table 1**), producing a collection of dose-response curves representing each individual ERCC in the study (**Figures 2 – 7, panel A**). ERCCs that were missing data for one or more concentrations in the RNA-Seq experiments were flagged as partially detected or undetected, and excluded from further analysis (**Figures 6 and 7, panel A**). The mid-point of each ERCC dose-response curve (average signal versus average relative abundance from the Latin square) was used to assess whether any particular ERCC was an outlier relative to the entire set of controls. The data were fit to an appropriate model for each platform (**Figures 2 – 7, panel B**).

**Figure 2.**
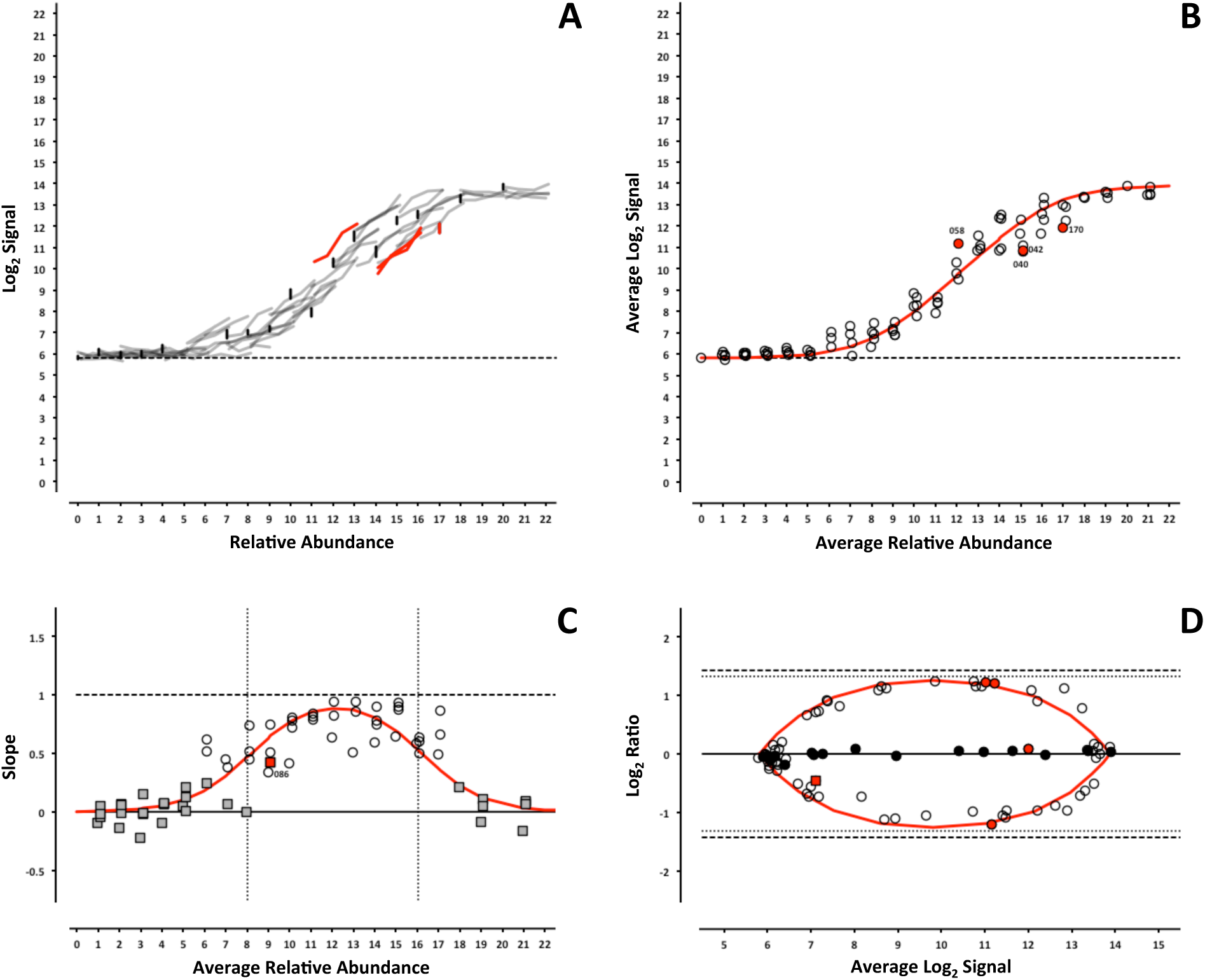
*ERCC signal response as a function of relative abundance in each of the four pools on the Illumina microarray platform*. In Panel A, each line represents an individual ERCC, where grey = titrated, black = 1-to-1, red = outlier, and dashed-line = background (average ERCC-00073). In Panel B, the centroid of each ERCC is plotted, where the red line corresponds to the fitted Langmuir model, open circles = within 99% CI, red circles = outliers, and dashed-line = background. In Panel C, the slope of each ERCC is plotted, where the red line corresponds to expected slope (first derivative of the Langmuir model), the vertical dotted lines correspond to the margins of the linear region (inflection points of the first derivative of the Langmuir model), the open circles = monotonic ERCCs *(fi* = 1), grey squares = non-monotonic, and red = outliers. Numbers in Panels B and C correspond to the last three digits of the Control ID in **Table 3**. In Panel D, each ERCC is represented on the Bland-Altman plot of Mix 1 vs Mix 3, where the red line corresponds to the ratio versus average intensity derived from the fitted Langmuir model, with outliers coded as in Panels B and C above.

**Table 3.** ERCC outliers grouped by performance criteria.

| Controls | Log2 Target<br>Relative<br>Abundance | Subpool | Response Curve Outliers |  |  |  |  | Non-monotonic |  |  |  |  |
| --- | --- | --- | --- | --- | --- | --- | --- | --- | --- | --- | --- | --- |
|  |  |  | AGL | ILM | NIAID | LifeTech | ILM HiSeq | AGL | ILM | NIAID | LifeTech | ILM HiSeq |
| ERCC-00156* | 1 | B |  |  |  | ○ |  |  |  |  | ○ |  |
| ERCC-00147 | 2 | C |  |  | ○ |  |  |  |  |  | ○ | ○ |
| ERCC-00148* | 2 | D | ○ [1,2] |  |  |  | ○ | ○ [1] |  |  | ○ | ○ |
| ERCC-00137 | 3 | B | ○ [2] |  |  |  |  | ○ [1] |  |  |  | ○ |
| ERCC-00143* | 3 | E | ○ [1] |  |  |  |  | ○ [1] |  |  |  |  |
| ERCC-00131* | 4 | D | ○ [1] |  | ○ |  |  | ○ [1,2] |  |  |  |  |
| ERCC-00134* | 4 | E | ○ [1] |  |  |  |  | ○ [1,2] |  |  |  |  |
| ERCC-00168* | 6 | E | ○ [1,2] |  |  |  | ○ | ○ [2] |  | ○ |  | ○ |
| ERCC-00095 | 8 | E |  |  |  | ○ |  |  |  | ○ |  |  |
| ERCC-00157 | 1 | C |  |  |  |  |  |  |  |  |  | ○ |
| ERCC-00158 | 1 | D |  |  |  |  |  | ○ [1] |  |  |  |  |
| ERCC-00160 | 1 | E |  |  |  |  |  | ○ [1] |  |  |  |  |
| ERCC-00145 | 2 | B |  |  |  |  |  | ○ [1] |  |  |  |  |
| ERCC-00150 | 2 | E |  |  |  |  |  | ○ [1] |  |  |  |  |
| ERCC-00138* | 3 | C |  |  |  |  |  | ○ [1,2] |  |  | ○ | ○ |
| ERCC-00142 | 3 | D |  |  |  |  |  | ○ [2] |  |  |  | ○ |
| ERCC-00128 | 4 | B |  |  |  |  |  | ○ [2] |  |  | ○ |  |
| ERCC-00111 | 6 | C |  |  |  |  |  |  |  |  |  | ○ |
| ERCC-00097 | 7 | B |  |  |  |  |  |  |  |  | ○ |  |
| ERCC-00098 | 7 | C |  |  |  |  |  |  |  | ○ | ○ |  |
| ERCC-00104 | 7 | E |  |  |  |  |  |  |  |  |  | ○ |
| ERCC-00086 | 8 | C |  |  |  |  |  |  | ○ |  |  | ○ |
| ERCC-00062* | 12 | C |  |  |  |  |  |  |  |  |  | ○ |
| ERCC-00046* | 13 | C |  |  |  |  |  |  |  |  |  | ○ |
| ERCC-00162 | 0 | A | ○ [1] |  |  |  |  |  |  |  |  |  |
| ERCC-00126 | 4 | A |  |  |  | ○ |  |  |  |  |  |  |
| ERCC-00113* | 5 | C | ○ [2] |  | ○ | ○ |  |  |  |  |  |  |
| ERCC-00117 | 5 | C |  |  | ○ |  |  |  |  |  |  |  |
| ERCC-00120 | 5 | D |  |  |  |  | ○ |  |  |  |  |  |
| ERCC-00109 | 6 | B |  |  | ○ | ○ | ○ |  |  |  |  |  |
| ERCC-00077 | 8 | A |  |  |  |  | ○ |  |  |  |  |  |
| ERCC-00081 | 9 | D |  |  |  |  | ○ |  |  |  |  |  |
| ERCC-00060 | 10 | A |  |  |  | ○ |  |  |  |  |  |  |
| ERCC-00075 | 10 | D |  |  |  | ○ |  |  |  |  |  |  |
| ERCC-00171 | 10 | B |  |  |  |  | ○ |  |  |  |  |  |
| ERCC-00054 | 11 | B |  |  |  |  | ○ |  |  |  |  |  |
| ERCC-00058* | 11 | D | ○ [1,2] | ○ | ○ | ○ |  |  |  |  |  |  |
| ERCC-00069 | 11 | E |  |  | ○ |  | ○ |  |  |  |  |  |
| ERCC-00044 | 12 | B |  |  |  | ○ | ○ |  |  |  |  |  |
| ERCC-00025 | 14 | A |  |  |  |  | ○ |  |  |  |  |  |
| ERCC-00028 | 14 | B |  |  |  |  | ○ |  |  |  |  |  |
| ERCC-00040 | 14 | C |  | ○ |  |  |  |  |  |  |  |  |
| ERCC-00042 | 14 | E |  | ○ |  | ○ |  |  |  |  |  |  |
| ERCC-00007 | 16 | D |  |  |  | ○ |  |  |  |  |  |  |
| ERCC-00022 | 16 | C | ○ [1,2] |  |  |  |  |  |  |  |  |  |
| ERCC-00170* | 16 | A | ○ [2] | ○ |  | ○ | ○ |  |  |  |  |  |
| ERCC-00023 | 18 | D | ○ [1,2] |  |  |  |  |  |  |  |  |  |
| ERCC-00002 | 20 | B |  |  |  |  | ○ |  |  |  |  |  |
| ERCC-00012 | 20 | A |  |  |  |  | ○ |  |  |  |  |  |
| ERCC-00024 | 20 | E | ○ [1,2] |  |  |  |  |  |  |  |  |  |

<sup>1</sup>Agilent 1-color data.
<sup>2</sup>Agilent 2-color data.
\*Discussed further in main text.
**Note:** The following analytes were incorrectly prepared as their antisense sequence and omitted from the data analysis: ERCC-00009, ERCC-00014, ERCC-00057, ERCC-00059, ERCC-00099, ERCC-00108, and ERCC-00116.

**Figure 3.**
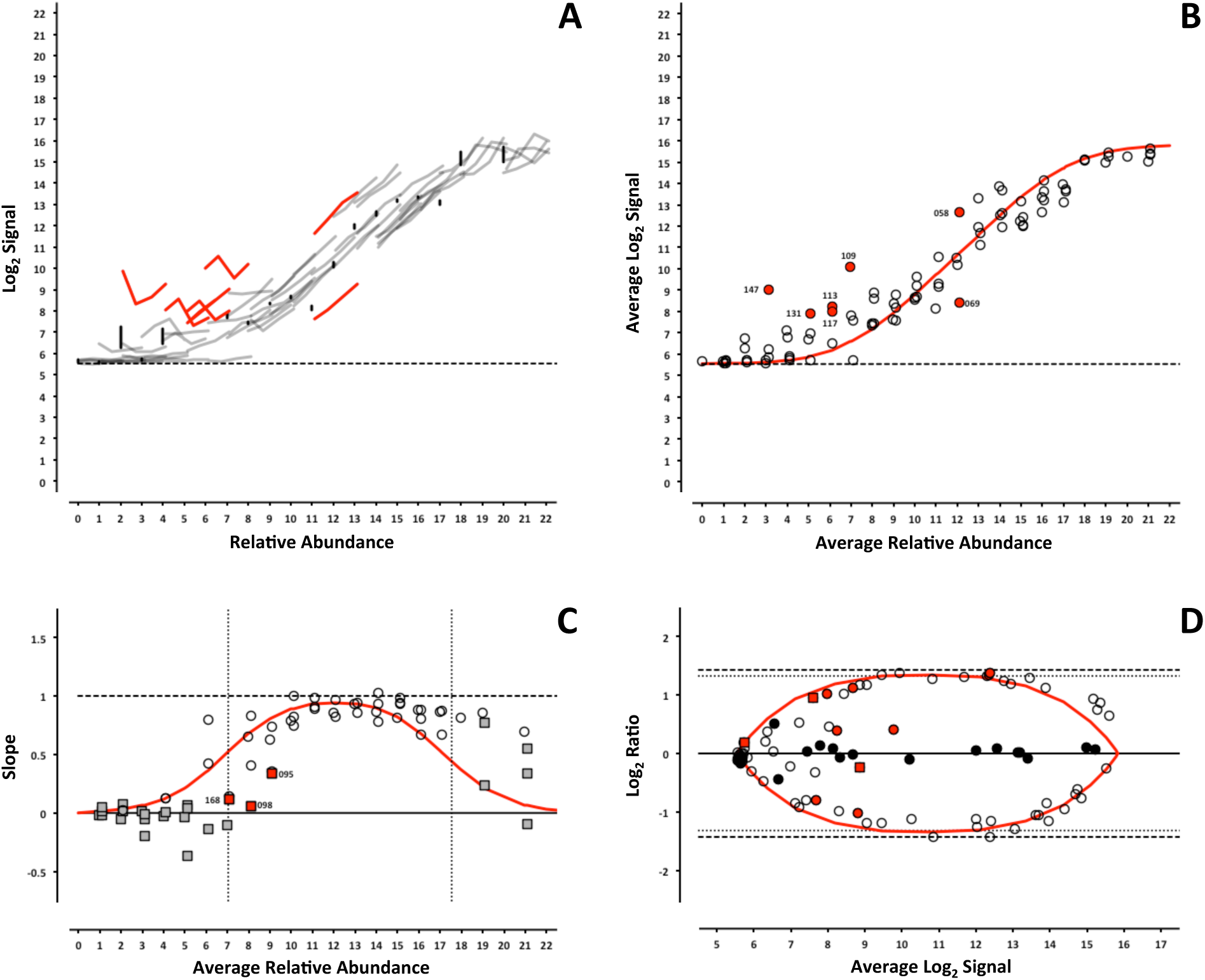
*ERCC signal response as a function of relative abundance in each of the four pools on the NIAID microarray platform*. See Figure 2 legend.

**Figure 4.**
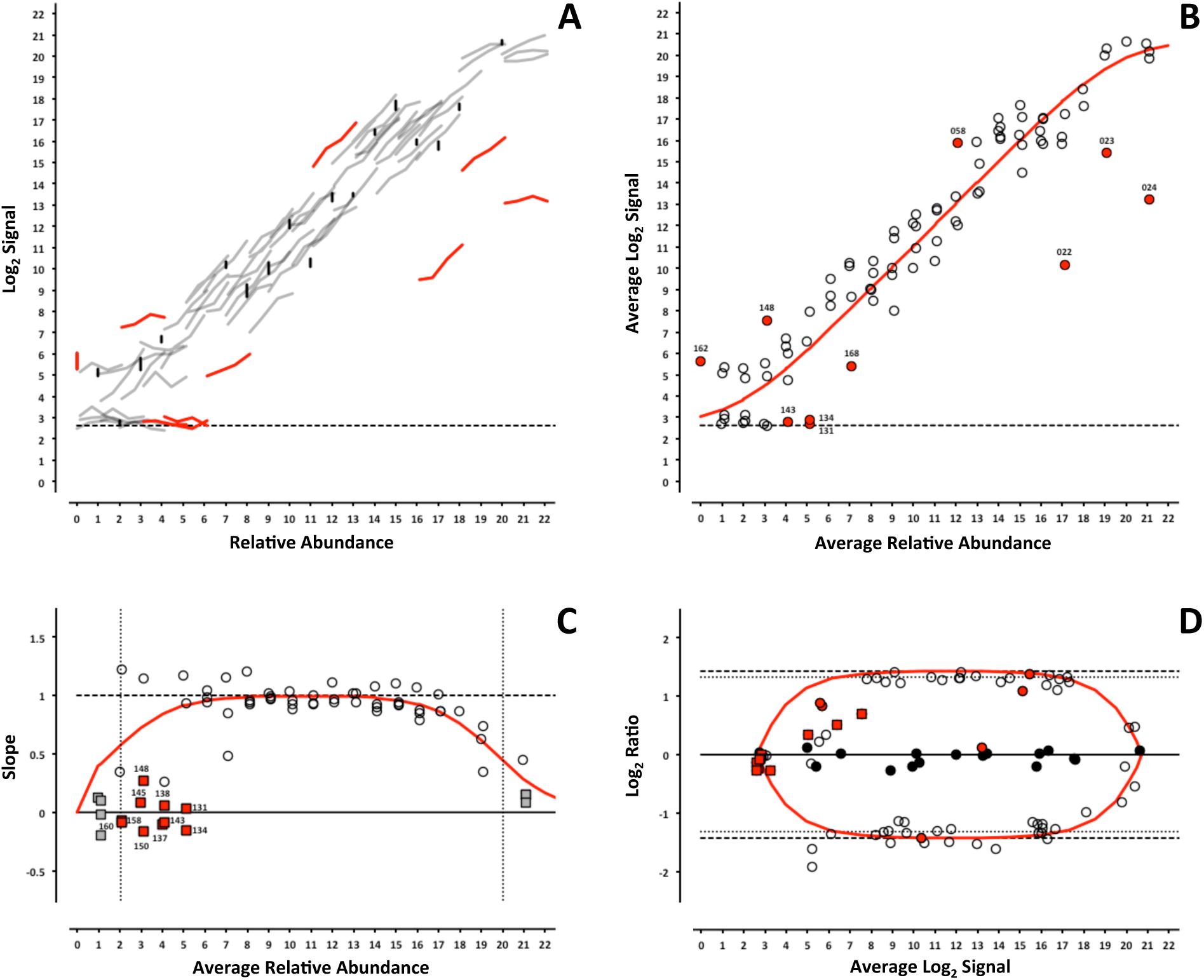
*ERCC signal response as a function of relative abundance in each of the four pools on the Agilent 1-color microarray platform*. See Figure 2 legend.

**Figure 5.**
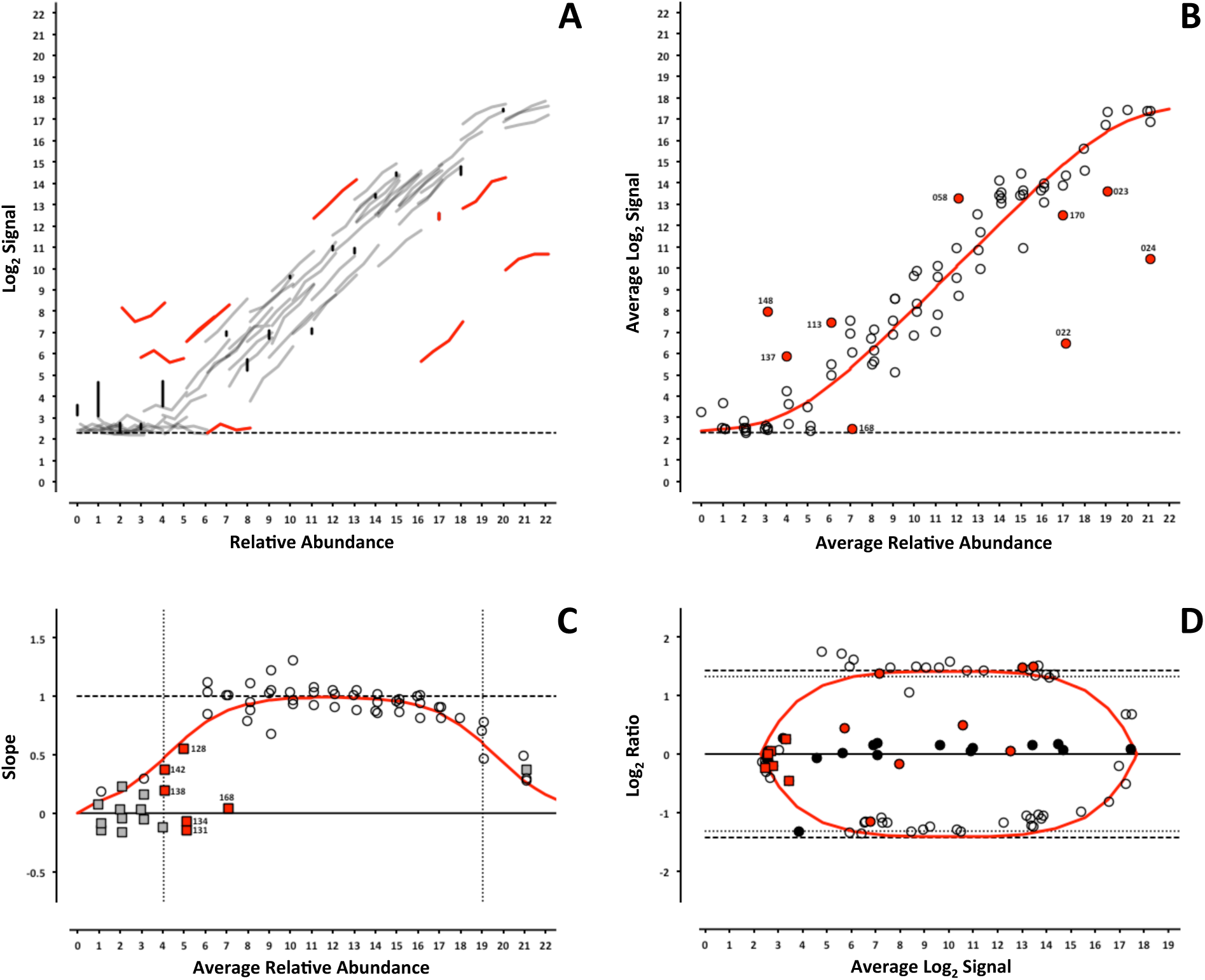
*ERCC signal response as a function of relative abundance in each of the four pools on the Agilent 2-color microarray platform*. See Figure 2 legend.

**Figure 6.**
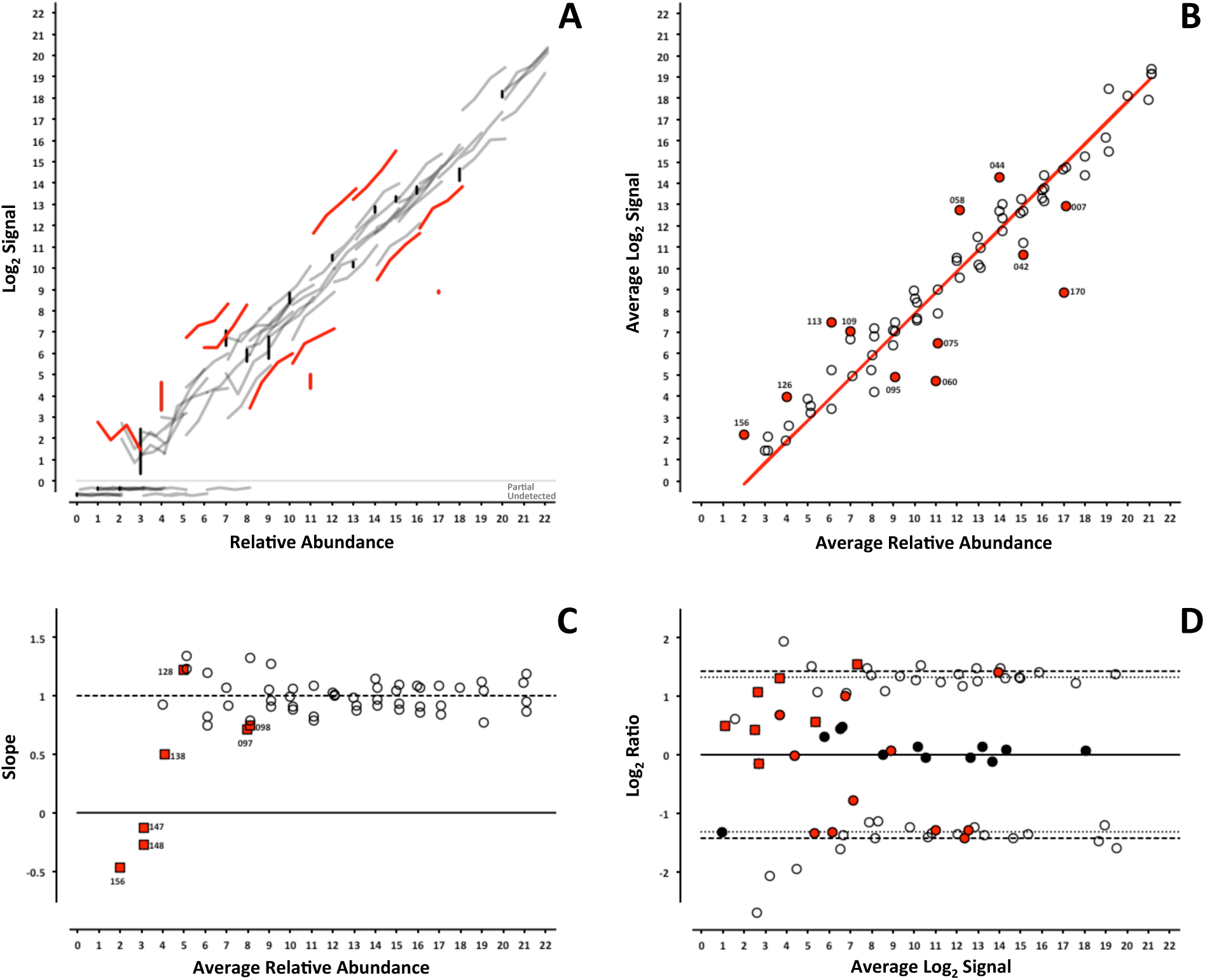
*ERCC signal response as a function of relative abundance in each of the four pools on the LifeTech NGS platform*. In Panel A, each line represents an individual ERCC, where grey = titrated, black = 1-to-1, and red = outlier. Partially detected and undetected ERCCs are included at the bottom to indicate their targeted relative abundance. In Panel B, the centroid of each ERCC is plotted, where the red line corresponds to the linear fitted model, open circles = within 99% CI, and red circles = outliers. In Panel C, the slope of each ERCC is plotted, where the open circles = monotonic ERCCs (*ρ* = 1), grey squares = non-monotonic, and red = outliers. Numbers in Panels B and C correspond to the last three digits of the Control ID in **Table 3**. In Panel D, each ERCC is represented on the Bland-Altman plot of Mix 1 vs Mix 3, with outliers coded as in Panels B and C above.

**Figure 7.**
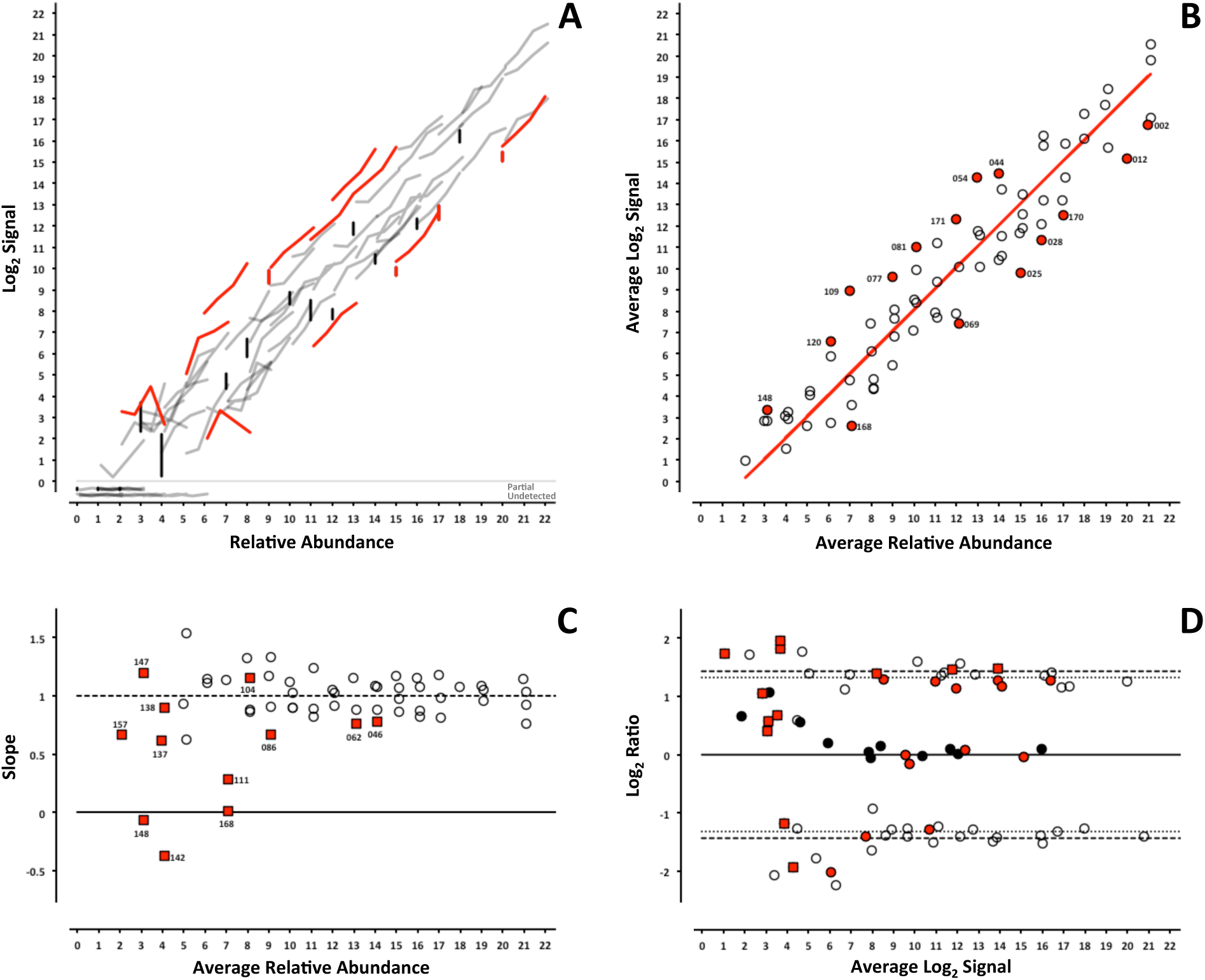
*ERCC signal response as a function of relative abundance in each of the four pools on the Illumina NGS platform*. See Figure 6 legend.

For the microarray experiments, a model using the Langmuir isotherm and was used [7, 8]. The dissociation constant, *K_d_*, was determined by fitting the data as follows:

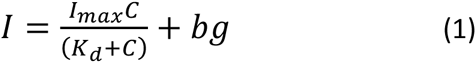

Where the maximal intensity of a feature at saturation, *I_max_*, and the background, *bg*, are experimentally derived from the average of the most abundant ERCC in each of the 4 pools and ERCC-00073, a component omitted from the pools, respectively. For the RNA-Seq experiments, a linear fit with a slope of 1 and fitted y-intercept was used as the model. For either model, ERCCs outside the 99% confidence interval (CI) were flagged as outliers (Figs 2 – 7, panel B) and compared across platforms to identify any ERCC-specific anomalies (**Table 3**).

With the exception of the ERCCs in the 1-to-1 subpool, the signal for each control should follow a strictly increasing monotonic function determined by the pool fraction of the Latin square design, 10% < 15% < 25% < 40%, (see **Fig. 1B**). This monotonicity was assessed with Spearman’s rho, *ρ*, where ERCCs with *ρ* < 1 were identified for comparison across platforms. In addition, the slope of each individual ERCC dose-response curve can be calculated and plotted as a function of the relative abundance, where the slope (*m* = 1) corresponds to an ideal dose-response. For the microarray data, the first derivative of the Langmuir function also provides us with a model of the expected slope and the inflection points allow us to demarcate a region of the dynamic range where we should expect a linear response (**Figs 2 – 5, panel C**) [9]. Non-monotonic ERCCs that fall within that portion of the dynamic range were also identified as outliers. For the RNA-Seq data, all non-monotonic ERCCs are flagged as outliers (**Figs 6 and 7, panel C**). One control, ERCC-00113, was an outlier on all platforms, with *ρ* = −0.2 for each. Closer inspection of the monotonic trend indicated that the least abundant target feature produced the highest signal in each case. This ERCC was more consistent with membership in subpool C, indicating a likely error in the preparation of the subpools. Therefore, **Figures 2 – 7** include this ERCC plotted as a component of subpool C.

**Table 3** includes all ERCC identified as outliers by the two criteria above and highlighted in **Figures 2 – 7** and specific controls discussed below are indicated with an asterisk (*). The majority of non-monotonic ERCCs in the microarray experiments occurred below the lower inflection point on the slope plots and those flagged for non-detection in the RNA-Seq experiments also appear in the lower range of the signal response curves. For these ERCCs, it is difficult to assess performance beyond their utility for defining the lower limits of the linear range, so these are not included in the outlier table.

There were nine controls that were outliers on at least one platform for each criteria. Six of those were outliers for both criteria on the same platform: ERCC-00156 on LifeTech; ERCC-00131, ERCC-00134 and ERCC-00143 on AGL-1; ERCC-00148 on AGL-1 and ILM HiSeq; and ERCC-00168 on AGL-2 and ILM HiSeq. All of these controls performed well on the majority of platforms.

Fifteen ERCCs were non-monotonic only. ERCC-00046 and ERCC-00062 were the most highly abundant outliers in this class. In both cases, the two lowest concentrations for each control produced nearly identical values where the lowest concentration is slightly higher. With the exception of ERCC-00138, all of these controls performed well on the majority of platforms.

There are 26 ERCCs that appear to be outliers with respect to the overall dose-response model that are still monotonic. For example, ERCC-00058 was the only control to be determined a response curve outlier on all microarray platforms and one RNA-Seq platform, however the observed slope on all platforms tested was greater than 0.9. ERCC-00170 was also flagged on every platform except the NIAID microarray, but was not evaluated for montonicity because it is in the 1-to-1 subpool.

Some of these results may be attributable to difficulties with accurately preparing large dynamic range pools with multiple controls, so that the actual concentration is different than the nominal abundance. The linear signal responses indicate the proper combinations of the subpools A – E were achieved for the Latin square design. Some of these outliers might also be the result of an RNA processing bias that may be analyte specific and proportional to abundance, for example poly-A enrichment [10].

### Intensity-dependent differential expression

For microarray data, an intensity-dependent bias is often visualized using an MA-plot; where M is the log2 transformation of the ratio of red and green fluorescence intensities in 2-channel data, and A is the log2 transformation of the average of the two [11]. This view has also been applied to two-condition single channel data, where M becomes the ratio of two different conditions, which is also referred to as a ratio-intensity plot (RI-plot) [12]. These comparative visualizations have been extended to sequencing data in the form of RA-plots, where the ratios and averages of integer count data form a characteristic pattern at the lower end of the signal range [13]. Each of these visualizations is a variation of a Bland-Altman plot (or difference plot), which is used here to visualize the ability to detect the nominal differences between two measurements [14]. A Bland-Altman plot of the ERCC components can be generated for any pairwise combination of Pools 12 – 15. One possible pairwise comparison, which produces fold-changes of 2.5 and 2.7 in both “up” and “down” directions (see **Fig 1B**) is shown in **Figures 2 – 7, panel D**. Additional pairwise comparisons are shown in **Supplemental Figures 1 – 6**.

For the microarray platforms, the discrimination between the target ratios is optimal near the middle of their dynamic range, and the ratios are “compressed” at both the lower and upper extremes. This constraint upon log_2_ ratios has been previously described [15]. The ratios converge towards unity at lower end due to background noise, which is additive, and contributes to both samples being compared. A similar compression is seen at high signal, where saturation dominates. We can also use **Equation 1** to derive the expected intensity ratios and average intensities for any fold-change of relative abundance. These fitted curves are also shown in **Figures 2 – 5, panel D**.

Signals in RNA-Seq are not subject to saturation (though high abundance transcripts can dominate the counting, and “crowd out” signals from lower abundance controls). As a consequence, the ratios do not compress at the upper end of the dynamic range. The RNA-Seq signals in this dataset are derived from counting technical replicates, where the variation can be characterized by a Poisson distribution [16]. In this case, “shot noise” dominates the signal at the low end, where counts might be added to either sample, and the ratios may deviate from target values in either direction **(Figs 6 and 7, panel D)**.

## Conclusions

The modified Latin square design provided for simultaneous evaluation of multiple controls with a minimal number of samples. While each individual ERCC was only tested over a small range of relative abundance, up to 4-fold for the ERCCs tested at multiple ratios and a single relative abundance value for the 1-to-1 components, in aggregate, they describe the overall measurement behavior of a platform.

The spread of the data indicates that differences in signals observed between different RNA species within the same sample may not accurately reflect the relative abundance between different RNA components of the same sample. Some of this dispersion may be due to the complexity of the pools used in these experiments where the distribution of target abundances described in Fig. 1A may not have been attained. For microarrays, probe designs for each ERCC target may also introduce some variability in signal between different ERCCs at the same relative abundance. For RNA-Seq, a non-uniform distribution of reads along different control sequences may also contribute to the variability [17].

The ERCCs did demonstrate that there is a linear region of the dynamic range of each platform where changes in abundance of a particular RNA transcript can produce a proportional change in signal. In this region, the ratios obtained with each platform approach the target ratios of the modified Latin square design. As a consequence, comparisons between samples for any particular RNA species can be expected to be accurate with respect to ratio-based measurements if they fall within this region. A pair of complex mixtures of RNA controls derived from NIST SRM 2374 designed to provide a set of ratios across a similar dynamic range is commercially available (Ambion™ ERCC ExFold RNA Spike-In Mixes). NIST has developed an R-based tool, the *erccdashboard*, to provide metrics and visualizations for these controls [18].

The ERCC RNA controls demonstrated utility in four different gene expression microarray platforms and two RNA-Seq platforms. Performance issues with any individual ERCC could be attributed to detection limitations or a target probe issue for particular platforms. The spike-in RNA controls were useful for establishing the dynamic range of relative abundance for a platform as well as delineating a reliable region where ratios can be measured accurately.

The composite testing scheme used in this study demonstrated that using well-designed pools of RNA controls provides measurement assurance for endogenous gene expression experiments. Pools of RNA controls from this study have been used as spike-ins for RNA-Seq experiments [19], and commercially available versions of these controls have been used for their intended purpose as quality controls [20–23]. These controls have also proven useful in product and method development due to their certified sequences and known concentrations [24–32]. Recently, they have become important in comparing transcriptomes between cell types in immunology [20, 32, 33], agriculture [34, 35], and other biology studies [21, 36–38], as well as key to understanding and accounting for the technical noise in single-cell sequencing experiments [39–42].

**Abbreviations**
ERCC: External RNA Controls Consortium
NIAID: National Institute of Allergy and Infectious Diseases
NIST: National Institute of Standards and Technology
SRM: Standard Refererence Material
PCR: Polymerase chain reaction
RNA-Seq: RNA sequencing
NGS: Next-generation sequencing
AGL-1: Agilent 1-color
AGL-2: Agilent 2-color
ILM-bead: Illumina bead array
ILM-ngs: Illumina next-generation sequencing
LifeTech: Life Technologies

## Competing Interests

JL is employed by Illumina Inc., manufacturer of one of the microarray platforms and one of the sequencing platforms used in this study. ABL is employed by Agilent Inc., manufacturer of one of the microarray platforms used in this study. These authors provided data. All data analysis was performed by NIST.

## Authors’ Contributions

ABL, JL, TGM, and SM designed the study. JM, SMJ-H, and SM developed the reference samples. SAM, JM, ABL, JL, TGM, and SQ acquired and processed the data. All authors participated in the preliminary analysis and interpretation. PSP developed metrics and visualizations and drafted the manuscript. All authors participated in the revision process and provided final approval.

## Acknowledgements

This research was supported in part by the Intramural Research Program of the NIH, NIAID.

## Disclaimer

Certain commercial equipment, instruments, or materials are identified in this paper in order to specify the experimental procedure adequately. Such identification is not intended to imply recommendation or endorsement by the National Institute of Standards and Technology (NIST), nor is it intended to imply that the materials or equipment identified are necessarily the best available for the purpose.

---

**Supplemental Figure 1.**
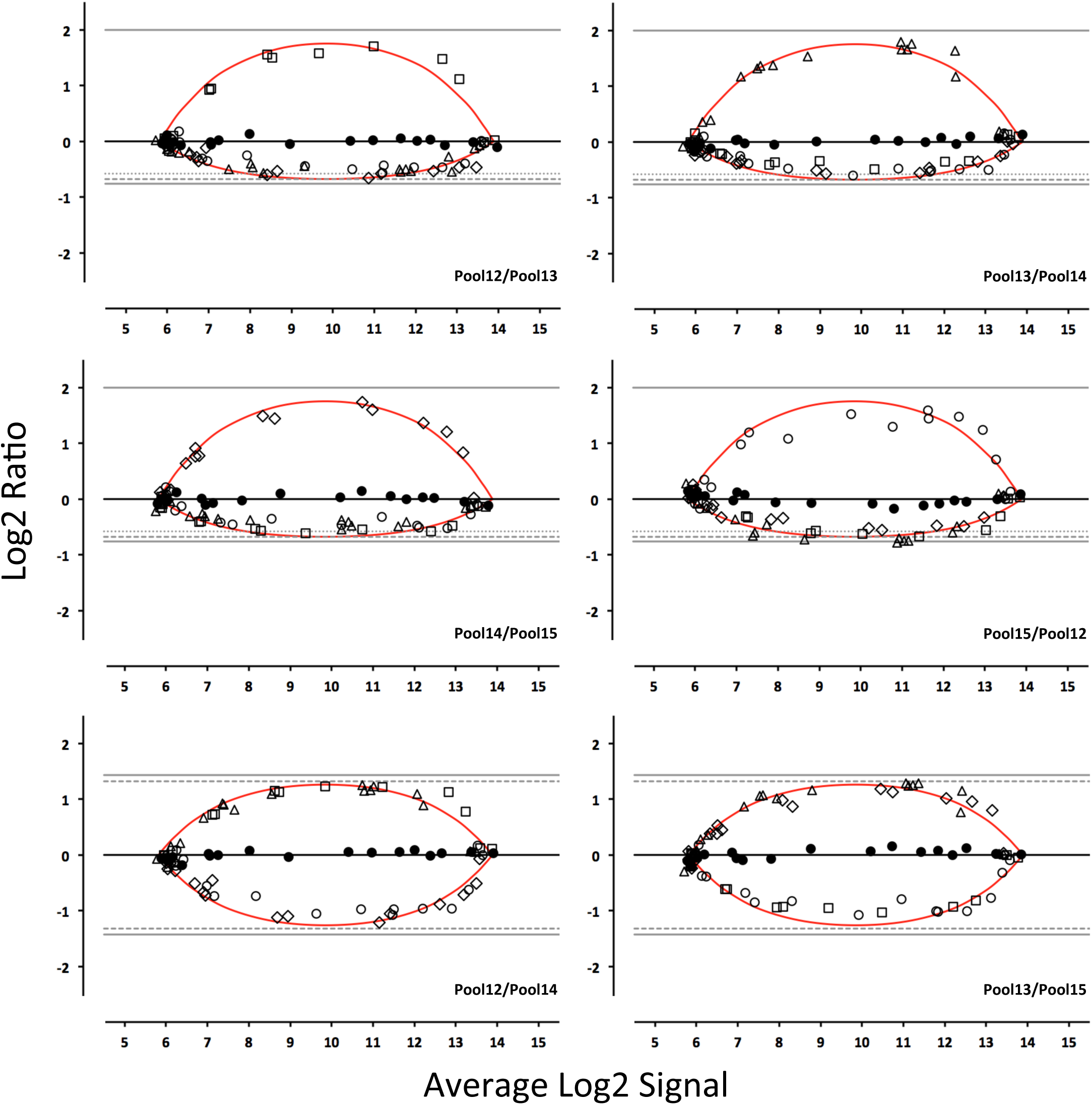
*Bland-Altman plot of each pair-wise pool comparison using the Illumina microarray platform*. Symbols correspond to pools A – E (see Fig. 1). Filled circles = A, open circles = B, open diamonds = C, open triangles = D, and open squares = E. The red line corresponds to the ratio versus average intensity derived from the fitted Langmuir model.

**Supplemental Figure 2.**
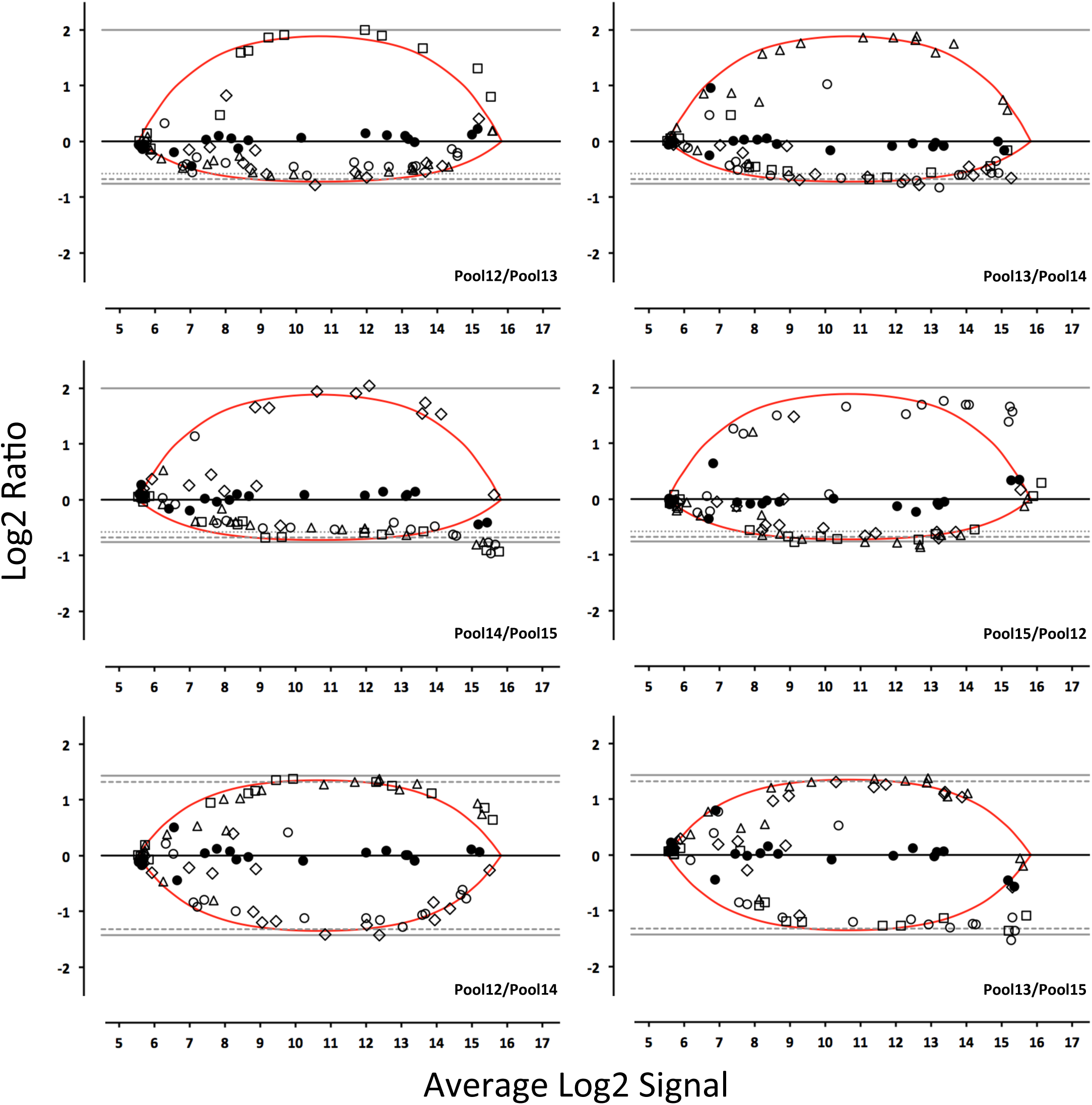
*Bland-Altman plot of each pair-wise pool comparison using the NIAID microarray platform*. See Supplemental Figure 1 legend.

**Supplemental Figure 3.**
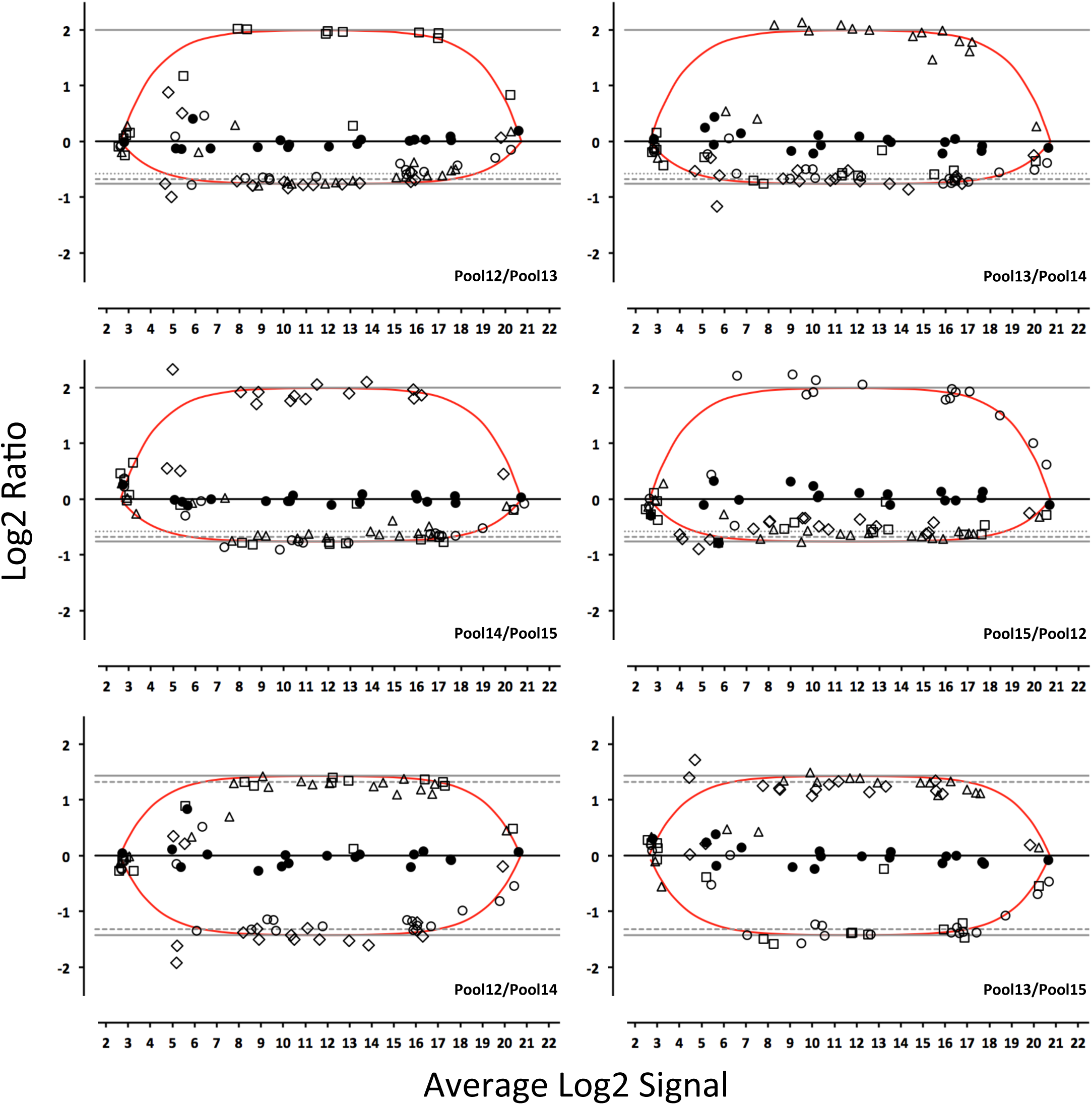
*Bland-Altman plot of each pair-wise pool comparison using the Agilent* 1 *color microarray platform*. See Supplemental Figure 1 legend.

**Supplemental Figure 4.**
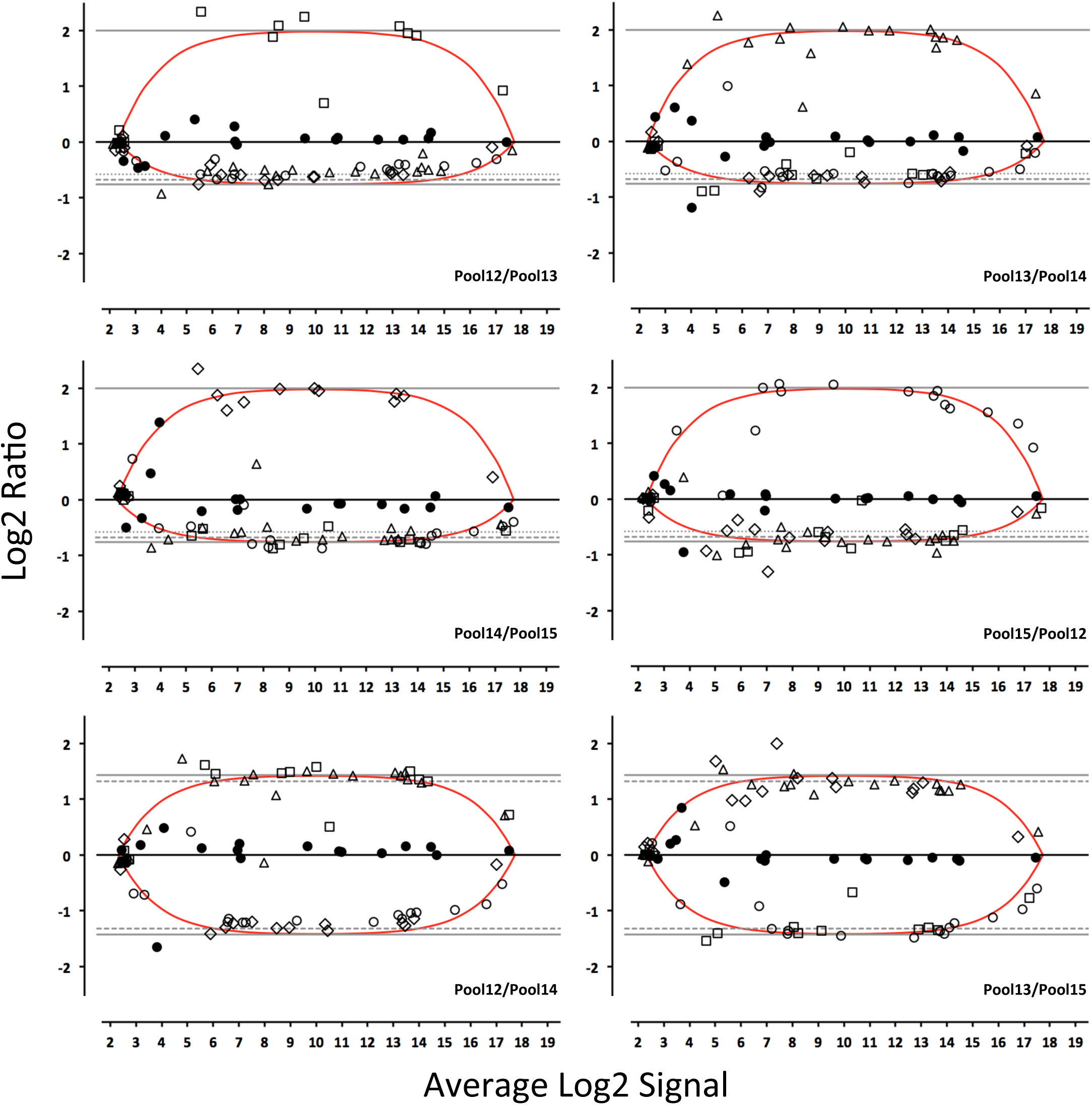
*Bland-Altman plot of each pair-wise pool comparison using the Agilent* 2 *color microarray platform*. See Supplemental Figure 1 legend.

**Supplemental Figure 5.**
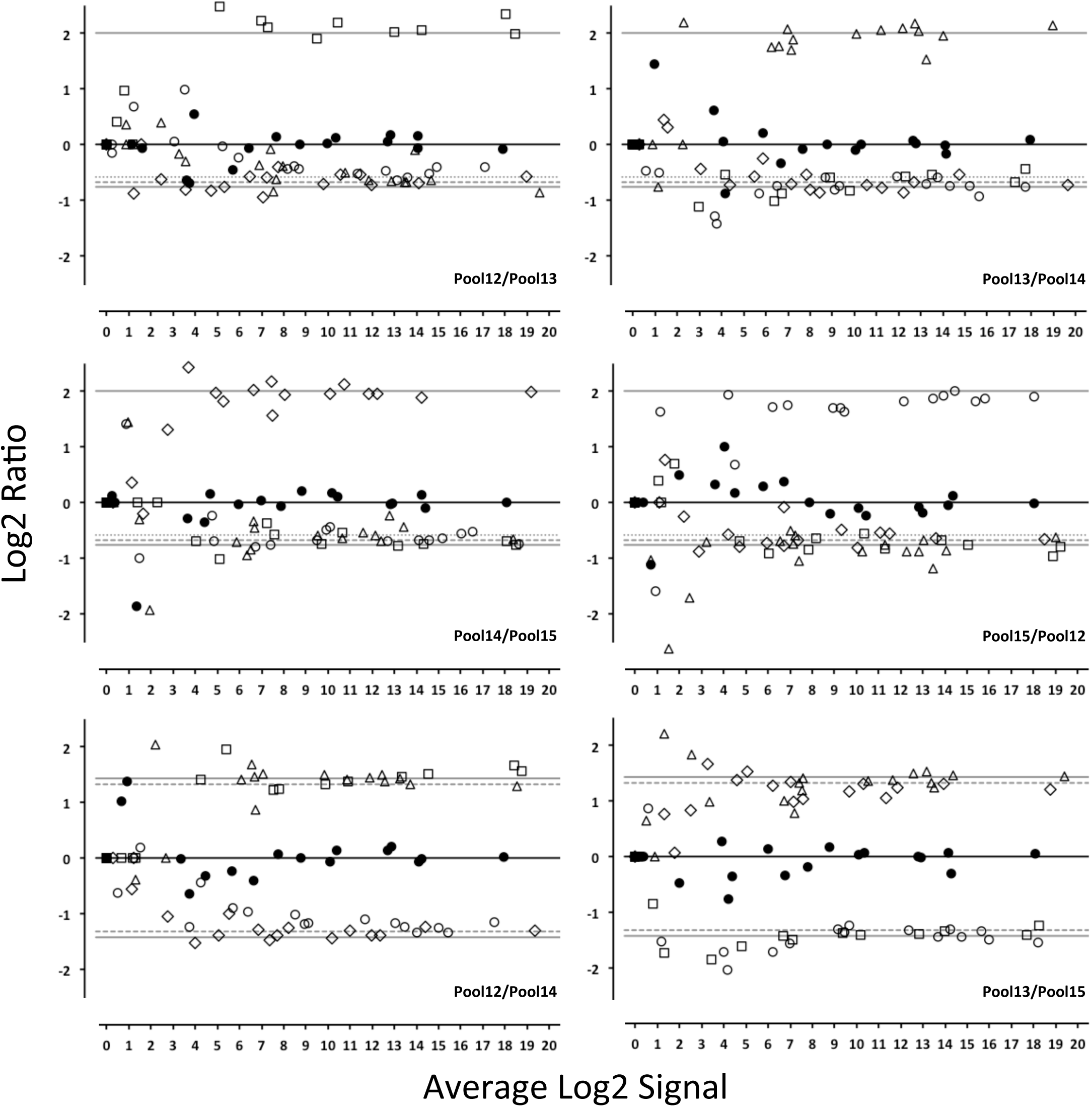
*Bland-Altman plot of each pair-wise pool comparison using the LifeTech NGS platform*. Symbols correspond to subpools A – E (see Fig. 1). Filled circles = A, open circles = B, open diamonds = C, open triangles = D, and open squares = E.

**Supplemental Figure 6.**
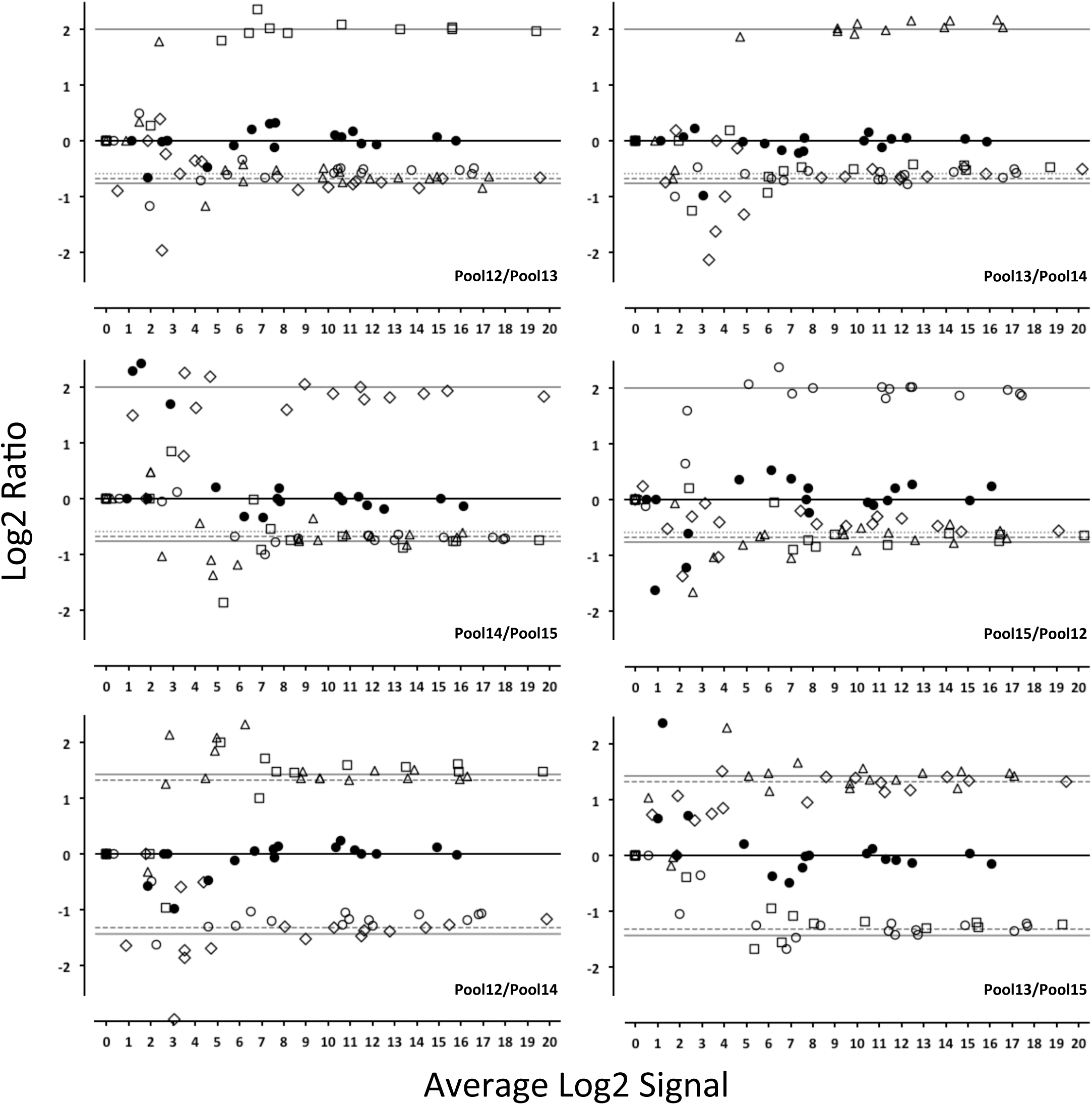
*Bland-Altman plot of each pair-wise pool comparison using the LifeTech NGS platform*. See Supplemental Figure 5 legend.

